# A small molecule reveals role of insulin receptor-insulin like growth factor-1 receptor heterodimers

**DOI:** 10.1101/2021.12.09.471780

**Authors:** Katie J Simmons, Chloe G Myers, Hema Viswambharan, Natalie J Haywood, Katherine Bridge, Samuel Turvey, Thomas Armstrong, Lydia Lunn, Paul J Meakin, Eva Clavane, David J Beech, Richard M Cubbon, Stephen B Wheatcroft, Martin J McPhillie, Colin WG Fishwick, Mark T Kearney

## Abstract

The insulin receptor and insulin like growth factor-1 receptor are heterodimers consisting of two extracellular α-subunits and two transmembrane β-subunits. IR αβ and IGF1R αβ hemi-receptors can heterodimerize to form hybrids composed of one IR αβ and one IGF1R αβ. Widely distributed in mammalian tissues, in contrast to IR and IGF1R the physiological function of hybrids is unclear. To identify tool compounds that inhibit hybrid formation we performed a high-throughput small molecule screen based on a homology model of hybrid structure. Our studies unveil a first in class quinoline-containing heterocyclic small molecule that reduces hybrids by >50% in human umbilical vein endothelial cells with no effect on IR or IGF1R expression. Downstream of IR and IGF1R our small molecule led to reduced expression of the negative regulatory p85α subunit of phosphatidylinositol 3-kinase, an increase in phosphorylation of its downstream target Akt and enhanced insulin and shear-induced phosphorylation of Akt. We show that hybrids have a role in human endothelial cell physiology distinct from IR and IGF1R.

## Introduction

Over the past four decades changes in human lifestyle have contributed to an explosion of obesity (Swinburn et al. 2011, Afshin et al. 2017) and its frequent sequelae insulin resistant type 2 diabetes mellitus (T2DM) (Guariguata et al. 2014). A poorly understood hallmark of obesity and T2DM is disruption of insulin signalling(Petersen et al. 2018, Roden et al. 2019), leading to dysregulation of cellular growth and nutrient handling (Samuel et al. 2016). The insulin receptor (IR) acts as a conduit for insulin-encoded information, which is transferred via a complex intracellular signalling network including the critical signalling nodes phosphatidylinositol 3-kinase (PI3-K) and the serine/threonine kinase Akt, to regulate cell metabolism(Taniguchi et al. 2006). During evolution the IR and insulin-like growth factor-1 receptor (IGF1R) diverged from a single receptor in invertebrates(Garofalo 2002, Tan et al. 2011), into a more complex system in mammals(Belfiore et al. 2009). Stimulation of IR or IGF1R initiates phosphorylation of IR substrate (IRS) proteins at multiple tyrosine residues(Taniguchi et al. 2006), phosphorylated IRS1 binds PI3-K initiating the conversion of the plasma lipid phosphatidylinositol 3,4,-bisphosphate to phosphatidylinositol 3,4,5-trisphosphate (PIP3) which activates the multifunctional serine-threonine kinase Akt(Manning et al. 2007).

In endothelial cells (EC), Akt activates the endothelial isoform of nitric oxide synthase (eNOS) by phosphorylation of Ser1177 (Dimmeler et al. 1999, Fulton et al. 1999). In humans and other mammals despite high structural homology and activation of similar downstream pathways the biological processes regulated by insulin and IGF1 are strikingly different(Cai et al. 2017). Consistent with this in EC we demonstrated that deletion of IR reduced(Duncan et al. 2007, Sukumar et al. 2013), whereas deletion of IGF1R increased basal ^Ser1177^peNOS and insulin-mediated ^ser 1177^peNOS (Abbas et al. 2011). More recently we showed that increasing IR in EC enhances insulin-mediated ^ser473^pAkt but blunts insulin-mediated ^Ser 1177^peNOS (Viswambharan et al. 2017), whereas increased IGF1R reduces basal ^ser1177^peNOS and insulin-mediated ^ser1177^peNOS(Imrie et al. 2012).

The tyrosine kinases IR and IGF1R are homodimers consisting of two extracellular α-subunits and two transmembrane spanning β-subunits(De Meyts et al. 2002). IR and IGF1R can heterodimerize to form hybrid receptors composed of one IR αβ complex and one IGF1R αβ complex(Soos et al. 1993). In skeletal muscle, fat and the heart, hybrid expression has been shown to exceed that of IGF1R and IR(Bailyes et al. 1997). While the role of hybrids in human physiology is undefined there is a clear association with increased hybrids and situations of metabolic stress including: T2DM (Federici et al. 1996, Federici et al. 1997), obesity, hyperinsulinemia(Federici et al. 1998), insulin resistance(Valensise et al. 1996) and hyperglycaemia(Federici et al. 1999). Hybrids are not amenable to genetic manipulation so development of small molecules to inhibit hybrid formation is an attractive approach to examine their physiological role. Here we describe the rational, structure-based discovery of a small molecule that inhibits formation of hybrids in human cells. This small molecule reveals a unique role for hybrids in cell biology answering a long unresolved question in human biology.

## Results and Discussion

### Virtual screening to identify small-molecule inhibitors of insulin receptor-insulin like growth factor receptor hybrids

As there is no published structure for hybrids, we generated a homology model of the hybrid receptor ectodomain (Figure 1A) based on the IR ectodomain dimer described by Croll et al(Croll et al. 2016). We examined this hybrid homology model using the KFC2 protein-protein interaction hotspot prediction server(Zhu et al. 2011) to identify key regions involved in the IR/IGF1R protein-protein interaction (**Figure 1B**). Each of three hotspots identified from the KFC2 webserver was evaluated to determine their suitability for docking studies (**Figure 1C-E**). We selected a hotspot covering amino acids 400-570 in IR for virtual high-throughput screening (**Figure 1E**). This region had the lowest degree of sequence identity between IR and IGF1R thereby affording us the greatest likelihood of identifying a selective modulator of hybrid formation. A library of commercially available small molecules was then prepared using the ZINC database of drug-like molecules(Sterling et al. 2015) and the OpenEye conformer generation software to generate a low energy conformer for each entry. Glide high throughput virtual screening mode(Halgren et al. 2004) mounted on the ARC3 advanced research computing cluster at the University of Leeds, was used to predict the binding pose of each ligand. A subset of the 100,000 top scoring molecules was rescored using GLIDE SP, before rescoring using a separate function-AutoDock Vina(Trott et al. 2010). The best scoring molecules were further examined using our in house *de novo* design program SPROUT(Law et al. 2003) searching for specific H-bonding, van der Waals and hydrophobic interactions with IR, and steric clashes with IGF1R, which could potentially prevent hybrid formation (**Figure 1F**).

**Figure 1.**
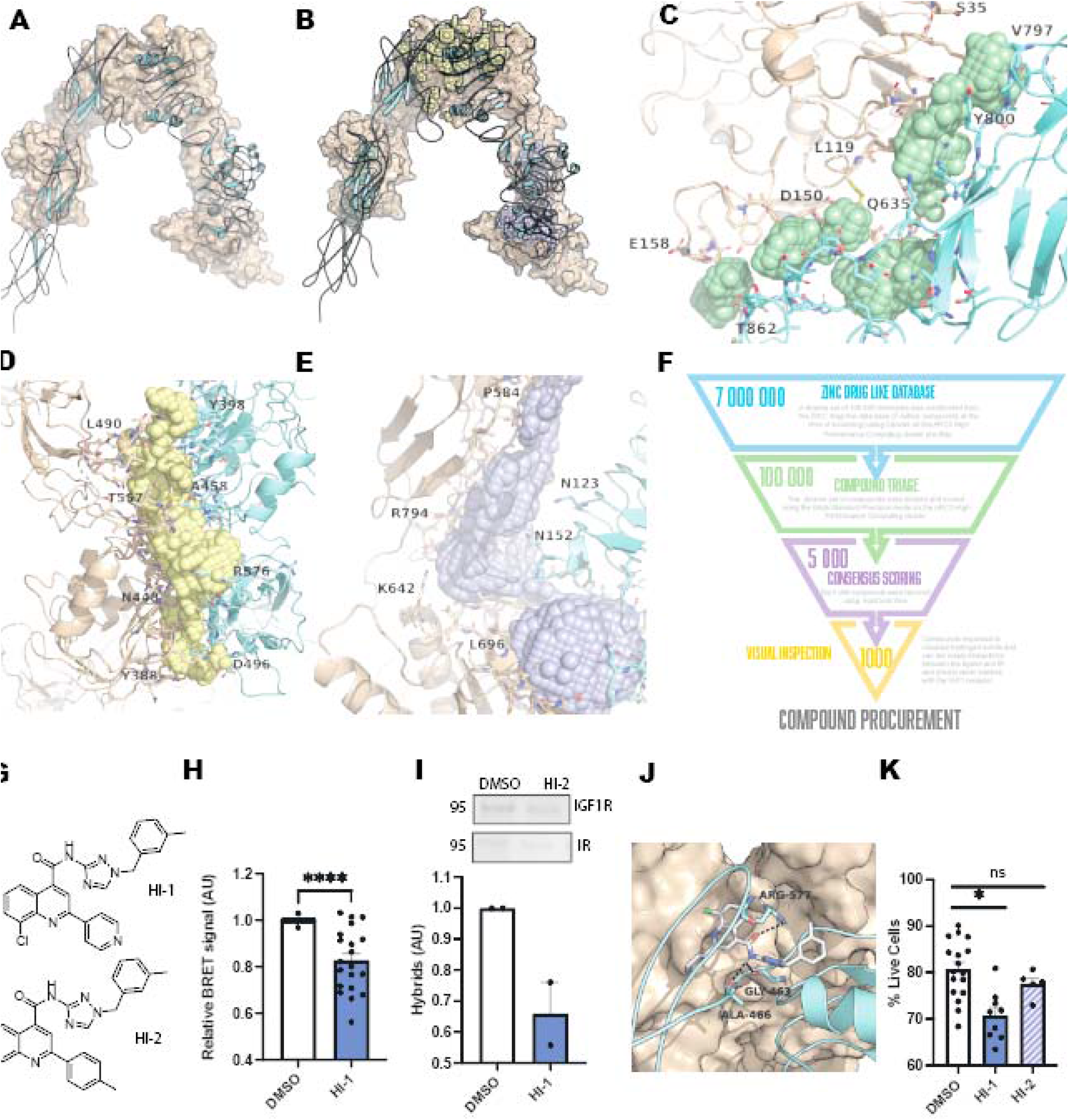
**A-** Homology model of the IR-IGF1R hybrid receptor produced using the I-TASSER webserver and the IR ectodomain dimer structure (PDB ID 4ZXB). The IR monomer is shown as a wheat surface and the IGF1R monomer is shown as cyan ribbons. Image generated using Pymol **B-** Hotspots identified as critical for IR-IGF1R hybrid receptor formation using the KFC2 webserver. The IR monomer is shown as a wheat surface and the IGF1R monomer is shown as cyan ribbons. Hotspot 1 is shown as green spheres, hotspot 2 is shown as yellow spheres and hotspot 3 is shown as lilac spheres. Image generated using Pymol **C-** Hotspot 1 (green spheres) covers residues 35-158 in the IR (wheat ribbons) and residues 635-862 in the IGF1R (cyans ribbons) A number of residues involved in the interaction are highlighted as wheat/cyan sticks. Image generated using Pymol **D-** Hotspot 2 (yellow spheres) covers residues 388-557 in the IR (wheat ribbons) and residues 398-576 in the IGF1R (cyans ribbons) A number of residues involved in the interaction are highlighted as wheat/cyan sticks. Image generated using Pymol **E-** Hotspot 3 (lilac spheres) covers residues 584-794 in the IR (wheat ribbons) and residues 123-152 in the IGF1R (cyans ribbons) A number of residues involved in the interaction are highlighted as wheat/cyan sticks. Image generated using Pymol **F-** Overview of the virtual high-throughput screening cascade using to identify potential inhibitors of hybrid formation. A diverse set of 100 000 molecules was constructed from the ZINC drug-like data base, the top scoring compounds using the Glide SP scoring function were then rescored using the AutoDock Vina scoring function, before the top 1000 compounds were visually inspected to visualise hydrogen bonds and van der waals interactions between the ligand and IR and predicted steric clashes with the IGF1R **G-** Structures of compounds **HI1** and **HI2** **H-** Compound **HI1** from the virtual high-throughput screen significantly reduced hybrid formation using a BRET assay in HEK293 cells compared to vehicle control (250 µM, 24h treatment, n=8,3 p= <0.001) **I-** Compound **HI1** from the virtual high-throughput screen reduced hybrid formation in HUVECs compared to vehicle control (250 µM, 24h treatment, n=3,2) **J-** Predicted binding mode of **HI1** to IR (wheat surface), making key hydrogen bonding interactions with Gly463, Ala466 and Arg577, but crucially makes a steric clash with Arg450 and Asn451 of IGF1R (cyan ribbons)-specifically preventing hybrid formation. A number of residues involved in the interaction are highlighted as wheat/cyan sticks. Image generated using Pymol **K-** Analysis of compound toxicity in HEK293 cells for the HI series showed that **HI1** significantly reduced cell viablilty, whereas there was no effect of **HI2** compared to vehicle control (cells treated for 24 hours, 100 uM compound, n=5,3 p=0.0013 for **HI1** and 0.2315 for **HI2**)

### To examine the effect of potentially bioactive molecules on hybrid formation we developed a Bioluminescence Resonance Energy Transfer (BRET) assay

HEK cells were co-transfected with cDNAs coding for IR-Rluc and IGF1R-YFP(Bacart et al. 2008). This results in the biosynthesis of three populations of receptors: (IR-Rluc)_2_ homodimers, (IGF1R-YFP)_2_ homodimers, and IR-Rluc/IGF1R-YFP heterodimers (**Figure S1**). Since only hybrids can produce a BRET signal, this method allows direct study of hybrids without having to separate them from homodimers of each type. To validate our assay, we showed increased BRET signal when cells were treated with IGF1 (100 nM) compared to basal levels, as described by Issad et. al. (Blanquart et al. 2006) (**Figure S2**).

We purchased a total of 42 compounds identified from a virtual high-throughput screen of seven million compounds (**Table S1**) from commercially available libraries and subjected them to an LCMS screen to determine compound purity and integrity. The BRET assay was used to find a potential candidate inhibitor of hybrid formation. This first compound we denoted as hybrid inhibitor-1 (**HI1**) (**Figure 1G**) reduced hybrids in the BRET assay (Figure 1H) and also in human umbilical vein endothelial cells (HUVECs) quantified using western blotting (Figure 1I) when used at 250 *μ*M concentration. **HI1** is predicted to bind to IR in the hotspot described above. It is predicted to make key hydrogen bonding interactions with Gly463, Ala466 and Arg577, but crucially makes a steric clash with Arg450 and Asn451 of IGF1R-specifically preventing hybrid formation (**Figure 1J**).

A LIVE-DEAD Cell Viability Assay was used to examine compound toxicity in HEK293 cells (**Figure 1K**). Compound **HI1** had some slight toxicity compared to cells treated with vehicle control (DMSO), this was flagged for analogue development. We developed a series of analogues of **HI1** (**Table S2**) from both in-house synthesis (43-56, **Table S2**) and commercial sources (**57-63, Table S2**, Vitas-M laboratories). **HI2** showed no significant cell viability effects compared to vehicle control and was also more potent, reducing hybrid formation significantly at 100 *μ*M concentration. Due to its structural similarity to our HI series, we also tested the neurokinin 3 receptor antagonist, talnetant (**64**, GlaxoSmithKline). We used the ligand-based screening tool ROCS(Hawkins et al. 2007, Hawkins et al. 2010) (OpenEye Scientific) to identify compounds from our in-house MCCB library of 30000 ligands which matched the shape and electrostatic properties of the HI series, but with a different central pharmacophore (**65** & **66, Table S2**).

Compounds produced in-house were synthesised using the schemes described in **Schemes S1-S3**. Briefly, 3-nitro-1*H*-1,2,4-triazole is coupled with arylalkyl chlorides before reduction of the nitro group to give the free amine. A small molecule X-ray crystal structure of the nitro triazole intermediate (**Figure S4** and **Tables S4-9**) confirmed the regioselectivity of the N-alkylation reaction. Ring-expansion of substituted isatins using the Pfitzinger reaction gave the 4-quinolinecarboxylic acid derivatives which are then coupled to the amino-triazole derivatives using the coupling reagent propylphosphonic anhydride (T3P). These compounds were screened in our BRET assay (**Figure S3**) and the most active of these compounds is henceforth described as **HI2** (**Figure 1H**).

### Preliminary absorption, distribution, metabolism, and excretion (ADME) analysis for HI series

Early-stage physicochemical profiling of **HI1** and analogues to ascertain compound metabolic stability and aqueous solubility *in vitro* (**Table S3**) showed that the HI series has a very short half-life in both human and murine liver microsomal cells. To progress this series towards *in vivo* analysis of hybrid function, new analogues of **HI2** would need to possess improved half-lifes. Compound aqueous solubility was measured at pH 7.4 in phosphate buffered saline (PBS) using the method described by Bellenie et. al.(Bellenie et al. 2020) Our initial hit, **HI1** possessed moderate solubility, but solubility was improved by replacement of the carbo-aromatic rings with heterocycles, correlating well with cLogP values for the series.

### Effect of HI2 on expression of hybrids phenotype in human umbilical vein endothelial cells

In HUVECs **HI2** reduced hybrid expression (Figure 2A) with no effect on total IR (Figure 2B) or IGF1R expression (Figure 2C). Expression of the genes encoding IR and IGF1R were unchanged in **HI2** treated HUVEC (Figure 2D). We went on to examine the effect of **HI2** on basal expression of molecules at critical signalling nodes in the insulin signalling pathway. After treatment with **HI2** there was a reduction in expression of the p85α subunit of PI3-K (Figure 2E) but no change in p110α (Figure 2F). **HI2** had no effect on basal expression of Akt (Figure 2G) but reduced expression of eNOS (Figure 2H). **HI2** led to an increase in ^Ser473^pAkt (Figure 2I) and ^Thr308^pAkt (Figure 2J), downstream of Akt, ^Ser1177^peNOS was unchanged even though total eNOS was reduced (Figure 2K). **HI2** had no effect on proliferation (Figure S4A) or tube forming (Figure S4B) in HUVECs. Consistent with previous reports showing PI3K/Akt signalling is important in endothelial sprouting(Davies et al. 2019) sprout number in bead sprouting assays was increased, (**Figure 2L&N**) with no change in sprout length (**Figure 2M&N**).

**Figure 2.**
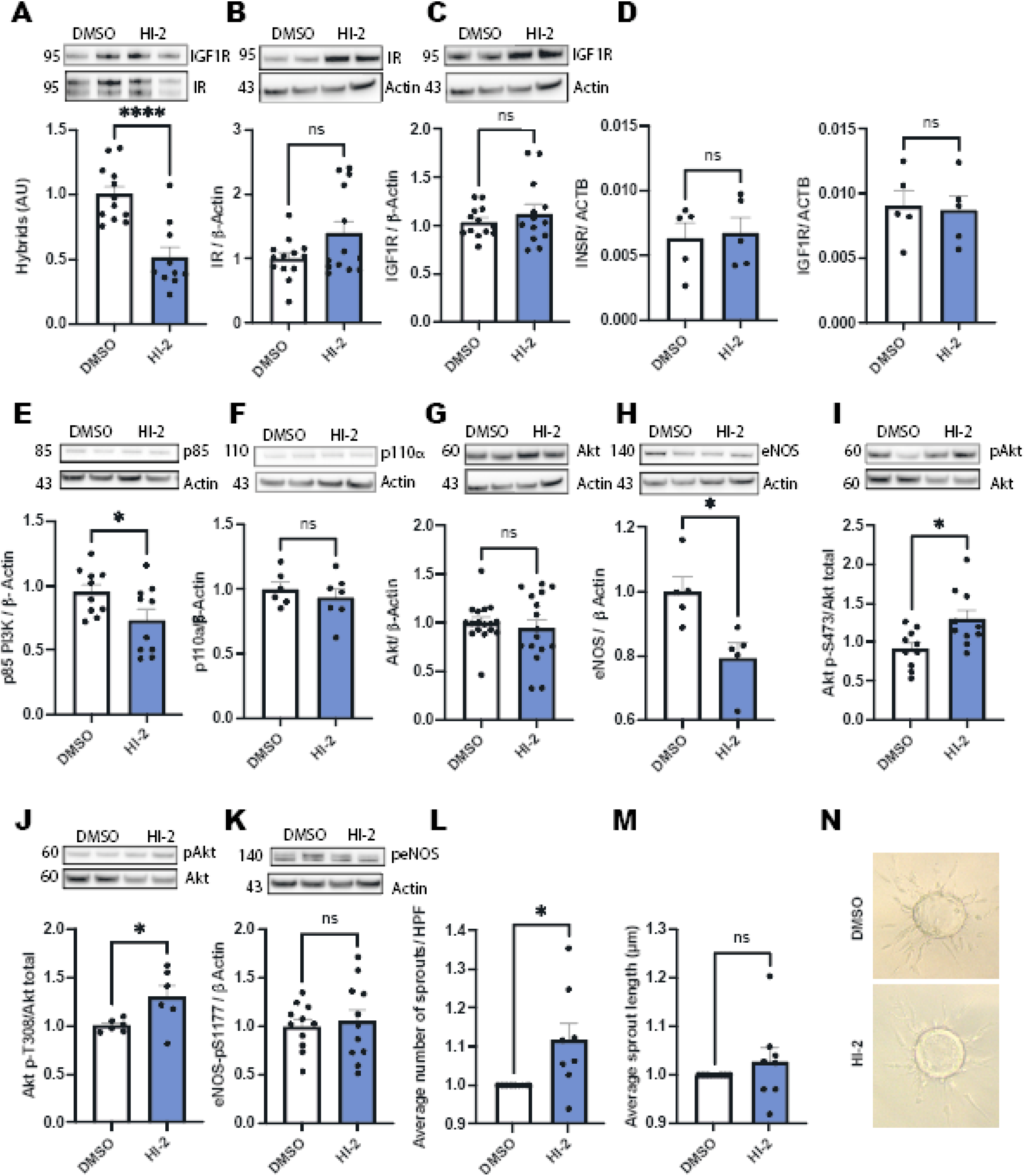
**A-** Quantification of hybrid formation in HUVECs treated with **HI2** compared to vehicle control (100 µM, 24h treatment, n=10 p= <0.001) **B-** Quantification of IR dimer formation in HUVECs treated with **HI2** compared to vehicle control (100 µM, 24h treatment, n=13 p= 0.07) **C-** Quantification of IGF1R dimer formation in HUVECs treated with **HI2** compared to vehicle control (100 µM, 24h treatment, n=13 p= 0.3867) **D-** Quantification of gene expression encoding IR and IGF1R in HUVECs treated with **HI2** compared to vehicle control (100 µM, 24h treatment, n=5 p= 0.8141 for IR and 0.8396 in IGF1R) **E-** Quantification of total p85 PI3K in HUVECs treated with **HI2** compared to vehicle control (100 µM, 24h treatment, n=10 p= 0.0417) **F-** Quantification of total p110α PI3K in HUVECs treated with **HI2** compared to vehicle control (100 µM, 24h treatment, n=6 p= 0.4585) **G-** Quantification of total Akt in HUVECs treated with **HI2** compared to vehicle control (100 µM, 24h treatment, n=16 p= 0.5581) **H-** Quantification of total eNOS in HUVECs treated with **HI2** compared to vehicle control (100 µM, 24h treatment, n=5 p= 0.0115) **I-** Quantification of ^Ser473^pAkt in HUVECs treated with **HI2** compared to vehicle control (100 µM, 24h treatment, n=10 p= 0.0142) **J-** Quantification of_Thr308_ pAkt in HUVECs treated with **HI2** compared to vehicle control (100 µM, 24h treatment, n=6 p= 0.0372) **K-** Quantification of_Ser1177_ peNOS in HUVECs treated with **HI2** compared to vehicle control (100 µM, 24h treatment, n=5 p= 0.0095) **L-** Quantification of bead sprout number in HUVECs treated with **HI2** normalised to vehicle control (100 µM, 24h treatment, n=4,2 p= 0.0415) **M-** Quantification of bead sprout length in HUVECs treated with **HI2** normalised to vehicle control (100 µM, 24h treatment, n=4,2 p= 0.4094) **N-** Representative images of bead sprouting experiments from vehicle control and **HI2**

### Impact of HI2 on endothelial cell insulin and shear induced signalling

We then examined the effect of **HI2** on insulin-mediated activation of Akt. Insulin induced ^Ser473^pAkt was increased across a range of insulin concentrations after treatment with **HI2** for 5 and 24 hours (**Figure 3A&C**). As described in landmark studies (Dimmeler et al. 1999, Fulton et al. 1999) Akt is activated by shear stress independent of ligand binding to cell surface receptors. We therefore examined the possibility that **HI2** would impact on shear induced pAkt. **HI2** had no effect on total Akt (**Figure 3B**) however it increased shear induced ^ser473^pAkt (**Figure 3D**).

**Figure 3.**
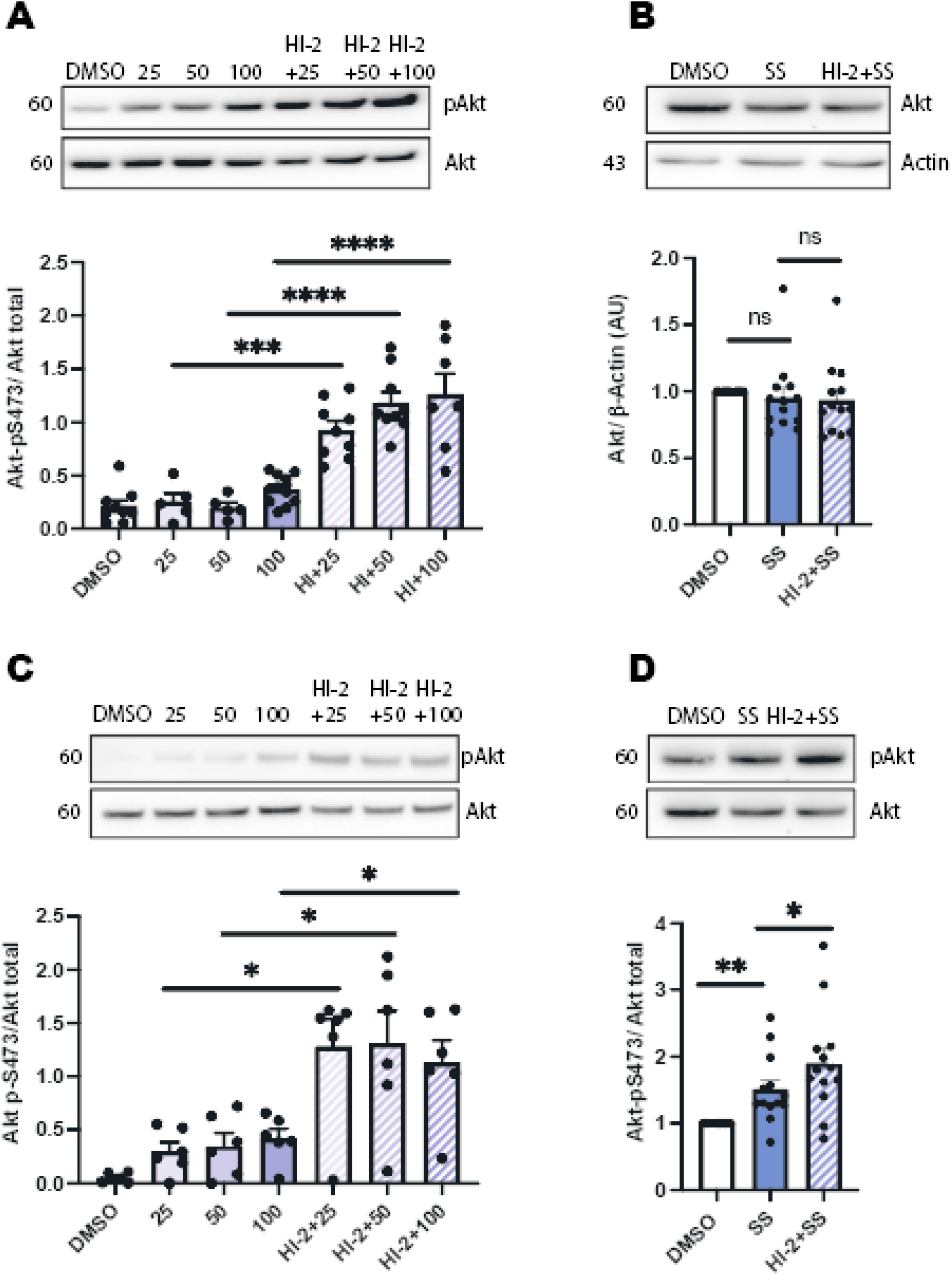
**A-** Quantification of insulin-induced ^Ser473^pAkt in HUVECs treated with **HI2** compared to vehicle control (100 µM **HI2** for 5 h, varying insulin concentration for 10 minutes, n= ≥5 p= <0.001) **B-** Quantification of shear-induced changes in total Akt in HUVECs treated with **HI2** normalised to vehicle control (100 µM, 24 h treatment, shear stress for 24 h, n=13 p= 0.8783) **C-** Quantification of insulin-induced ^Ser473^pAkt in HUVECs treated with **HI2** compared to vehicle control (100 µM **HI2** for 24 h, varying insulin concentration for 10 minutes, n=6 p=<0.05) **D-** Quantification of shear-induced _Ser473_pAkt in HUVECs treated with **HI2** normalised to vehicle control (100 µM, 24h treatment, shear stress for 24 h, n=15 p= 0.0235)

The IR and IGF1R are receptor tyrosine kinases which unlike most other receptor tyrosine kinases are preformed, covalently linked tetramers with two extracellular α subunits and two membrane-spanning, tyrosine kinase-containing β subunits. The physiological basis for the evolutionary preserved ability of IR and IGF1R to form hybrids has remained unclear as developing chemical approaches to disrupt protein–protein interactions is a significant challenge. Here we describe the identification, synthesis, and application of a first in class quinoline-containing heterocyclic small molecule inhibitor of IR/IGF1R hybrid formation. Following stimulation with insulin PI3-K, catalyses the conversion of the lipid phosphatidylinositol 3,4-bisphosphate to PIP3 which activates Akt(Engelman et al. 2006). PI3Ks are heterodimers consisting of a catalytic subunit (p110α, p110β, or p110δ) and regulatory subunit (p85α, p85β, p85γ, p50α, or p55α) (Engelman et al. 2006). The p110α isoform is ubiquitously expressed and essential for growth-factor signalling(Bi et al. 1999). Accumulating evidence suggests that monomeric free p85α acts as a negative regulator of PI3-K(Luo et al. 2005). Here we demonstrate that inhibition of hybrid formation increases basal, insulin and shear induced activation of this pathway.

## Materials and Methods

### Human Cell Line Cultures

HEK293 cells were cultured in Dulbecco⍰s Modified Eagle⍰s Medium (DMEM, Merck) containing 10% Fetal Calf Serum and 1% Antibiotic Antimycotic Solution (Sigma Aldrich). HUVECs were cultured in Endothelial Growth Medium-2 (ECGM-2, PromoCell) containing 10% Fetal Calf Serum and 1% Antibiotic Antimycotic Solution.

### Bioluminescense Resonance Energy Transfer Assay

Cells were seeded at a density of 200 000 cells per well in a six well culture plate and transfected with IR-Rluc only (control) or IR-Rluc and IGF1R-YPET (hybrids) using Lipofectamine 2000 as the transfection reagent (1 *μ*l of cDNA to 3 *μ*l of Lipofectamine 2000 per well) and incubated at 37 °C and 5% CO_2_. One day after transfection, cells were transferred into 96 well microplates (white, Poly-D-Lysine coated, Perkin Elmer, UK) at a density of 30,000 cells per well. On the following day (48 h after transfection), the cell culture medium was removed and replaced with media containing the compound of interest (100 *μ*M) or vehicle control (DMSO, 100 *μ*M). After a further 24 h cells were washed with 50 μl of phosphate-buffered saline solution (PBS, Sigma-Aldrich) prior to BRET measurements. BRET measurements were taken in PBS (final volume 50 μl) containing coelenterazine (stock solution (1 mM) in ethanol, final concentration 5 *μ*M). BRET measurements quantify Rluc light emission (485 nm) and YFP light emission (535 nm) using a Perkin Elmer microplate reader. BRET signal was calculated using the following equation:

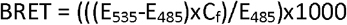

where C_f_= E_535_/E_485_ for the IR-Rluc only transfected HEK293 cells.

### LIVE/DEAD Cell Viability Assay

Quantification of cell viability was performed using the LIVE/DEAD Assay Kit (ThermoFisher), as per manufacturer’s protocol with minor modifications. Briefly, HEK293 cells were seeded at a density of 60 000 cell per well on a 96-well poly-L-lysine coated plate (Corning) and incubated overnight at 37 °C with 5% CO_2_. After incubation the desired compound was added to fresh media (DMEM, Merck) at a concentration of 100 µM before being added to the cells and incubated for 24 hours. After 24 hours, media containing the compound was removed and cells washed in PBS (200 µl). 20 µL of the 2 mM EthD-1 stock solution (Component B) was added to 10 mL of PBS to give a 4 µM EthD-1 solution. 5 µL of the supplied 4 mM calcein AM stock solution (Component A) was then added to the 10 mL EthD-1 solution. 100µL of the combined LIVE/DEAD® assay reagent was then added to each well and incubate for 45 min before quantification of percentage of red and green cells using the IncuCyte® ZOOM (Sartorius) system and its proprietary ZOOM 2016A software.

### HPLC Solubility Assay

Method was adapted from Bellenie et. al (Bellenie et al. 2020). Briefly, 10 µL of 10 mM DMSO stock solution was pipetted into a micro centrifuge tube (1.5 mL, Sarstedt) containing 990 µL of PBS buffer (pH 7.4, Sigma Aldrich) and mixed for 5 seconds on vortex mixer to create 100 µM solution with 1% DMSO. Following shaking of the suspension on a shaker at 500 rpm for 2 hours at room temperature (20 °C), it was separated by centrifugation (14000 rpm for 15 min). 200 µL of the supernatant was transferred to a 2 mL vial containing 50 µL of DMSO (Sigma Aldrich) and mixed for 5 seconds to avoid precipitation from the saturated solution. The concentration of the solubilized compound in solubility sample is measured by HPLC with UV detection using an external standard which was prepared by pipetting 10 µL of the same batch of compound DMSO stock used in solubility sample preparation to 990 µL of DMSO. The detail of the HPLC method is as following: chromatographic separation at 30°C is carried out over a 5 minute gradient elution method from 90:10 to 10:90 water:methanol (both modified with 0.1% formic acid) at a flow rate of 1.5 mL/min. Calibration curve is prepared by injecting 0.5, 2.5, and 5 µL of compound external standard. Compound solubility value is obtained by injecting 2 and 20 µL of compound solubility sample.

### Endothelial Cell Inhibitor Treatment

HUVECs were seeded at a density of 200 000 cells per well in a six well culture plate and incubated at 37 °C and 5% CO_2_. Cells were treated at 90-95% confluency. An aliquot of **HI2** (100 uM in DMSO) was added to ECGM-2 (PromoCell). Vehicle control treatments (DMSO, Sigma-Aldrich) wells were treated in an identical manner. Cells were incubated for 24 hours prior to lysis.

### Endothelial Cell Insulin Treatment

Cells were treated with **HI2** as described in Endothelial Cell Inhibitor Treatment protocol and incubated at 37 °C and 5% CO_2_. After 22 hours, the full-growth ECGM-2 medium was removed and replaced with M199 low-serum medium (2% FCS) containing **HI2**. At the end of the 24-hour treatment incubation period, insulin (10, 50, 100 nM, Sigma Aldrich) was added to the medium for 15 minutes at room temperature. To quench the reaction, ice-cold DPBS was added, and the cells were lysed as described in Endothelial Cell Lysis.

### Endothelial Cell Shear Stress

As we reported(Viswambharan et al. 2017) HUVECs were seeded at a density of 200 000 cells per well onto fibronectin-coated 6-well plates and incubated at 37 °C and 5% CO_2_. Once confluent, monolayers were placed onto an orbital rotating platform (Grant Instruments) housed inside the incubator. The radius of orbit of the orbital shaker was 10 mm, and the rotation rate was set to 210 rpm for 24 hours, which caused swirling of the culture medium over the cell surface volume. The movement of fluid because of orbital motion represents a free surface flow at the liquid–air interface at 12 dyne/cm

### Endothelial Cell Lysis

Cells were cultured in a six-well plate as described in the above methods. Prior to lysis, cells were washed twice in 2mL ice-cold PBS. A 100μL aliquot of Cell Extraction Buffer containing 1μL/mL protease inhibitor solution, was added to each well and left on ice for 5 minutes. The cells were scraped and collected and left on ice for 20 minutes, with intermittent mixing. A 20-minute centrifugation step at 4°C and 17,000xg was performed, collecting the supernatant into fresh receptacles. Protein quantification was performed using the Pierce ™ BCA Protein Assay kit (Thermo Scientific) as per manufacturer’s protocol.

### Immunoprecipitation

A solution containing 300μL of binding buffer (100 mM NaCl; 10 mM MgSO_4_; 100 mM HEPES; 0.025% Tween-20; pH 7.8) containing 1μL/mL protease and phosphatase inhibitors, and 30μL Protein G Agarose beads (Sigma-Aldrich) was combined with 3μL of primary antibody and 150 μg/μL cell lysate. Samples were incubated at 4°C overnight, for a minimum of 16-18 hours, on a medium speed MACs rotator. Upon completion, samples were centrifuged for one minute at 500xg, collecting the supernatant into a separate fresh receptacle. The samples were then washed in 1mL of PBS-T (0.05% Tween-20) containing 1μL/mL protease and phosphatase inhibitors, discarding the waste following each centrifugation and repeated six times in total. SDS NuPAGE™ LDS (4X) and Reducing Agent (10X) buffers (Invitrogen) were added to the beads. Samples were stored at -40°C until loading. Prior to loading, samples were boiled at 95°C for five minutes and centrifuged for one minute at 1000xg to pellet the beads. The supernatant was then used for immunoblotting.

### Immunoblotting

Non-immunoprecipitated protein samples (150 µg/µL) were also combined with SDS NuPAGE™ LDS (4X) and Reducing Agent (10X) buffers and boiled at 95°C for 5 minutes. Samples were placed on ice prior to loading. Samples were loaded onto NuPAGE™ 4-12% Bis-Tris Protein gels (Invitrogen) against a Colour Prestained Protein Standard (Broad Range 10-250 kDa, New England BioLabs) and resolved by electrophoresis for 70 minutes at 150-180V. Samples were transferred onto nitrocellulose or PVDF membranes using the TransBlot® Turbo™ Transfer system (Bio-Rad, USA) for 15 minutes at 20V. Non-specific sites on the membrane were blocked for 1 hour in 5% milk buffer (5% in 20 mM Tris pH 7.5; 145 mM NaCl; 0.05% Tween-20) at room temperature. Target proteins were labelled using their respective antibody (refer to **Table S1**0 for summary of antibodies used) in 5% Bovine Serum Albumin (BSA, Sigma-Aldrich) overnight at 4°C. The following morning, the membrane was further incubated with horse radish peroxidase (HRP) anti-mouse or anti-rabbit secondary antibodies (1:5,000) in 5% milk buffer for 1 hour at room temperature. The blot was then visualised using Immobilon® Western Chemiluminescent HRP Substrate reagents (Millipore) using the G:Box (SynGene) system.

### Real-Time Polymerase Chain Reaction

Confluent monolayers of HUVECs were lysed by addition of 0.05% trypsin (Invitrogen) and neutralized using ECGM-2 (PromoCell). Two wells of the 6-well plate, sharing the same compound or vehicle-treatment were combined and centrifuged at 1000xg for 8 minutes at 4°C. The cells were resuspended in DPBS (PromoCell) and centrifuged once more under the same conditions. The pelleted cells were dried and stored at -80°C until use. RNA extraction was performed using the Monarch Total RNA Miniprep Kit (NEB), according to manufacturer’s protocol. The resultant RNA samples were placed on ice for duration of the experiment. RNA quantitation was performed using the Nanodrop DS-11 FX+ Spectrophotometer/Fluorometer (DeNOVIX) system by addition of 2 μL of RNA sample to the crystal plate. Data recorded on the device was collected and exported into Microsoft Excel for further data transformation. Reverse transcription of RNA to cDNA was performed by combining 500 ng/ mL of the RNA samples with the LunaScript™ RT SuperMix (5X) (NEB) in 20 μL aliquots. cDNA samples were diluted in Nuclease-free water (NEB) before use in qPCR reactions. The qPCR samples were prepared by combining the cDNA samples with iTaq™ Universal SYBR® Green Supermix (BioRad) and analysed by the LightCycler® 96 system (Roche) using the following primers: IGF1R - PrimePCR™ SYBR® Green Assay: IGF1R, Human (Bio-Rad, USA) INSR - PrimePCR™ SYBR® Green Assay: INSR, Human (Bi-Rad, USA) and ACTB - PrimePCR™ SYBR® Green Assay: ACTB, Human (Bio-Rad, USA). Resultant data was analysed using the proprietary LightCycler® Software 4.1 and Microsoft Excel. The cycles to threshold” (cT) was measured for each well, the average of triplicate readings for each sample taken, normalised to ACTB.

### Bead Sprouting Assay

Method was adapted from Nakatsu et al. (Au - Nakatsu et al. 2007) Briefly, HUVECs were incubated at 37 °C and 5% CO_2_. in EGM-2 (Lonza) for 24 hrs prior to plating. Cytodex-3 Beads (Amersham, 10 mg/ml) were coated with HUVECs and incubated for 18 hours in T25 flask, before being suspended in a fibrinogen (2 mg/ ml, Sigma-Aldrich) /aprotinin (4 U/ml, Sigma-Aldrich) solution containing 5 ng/ml VEGF and FGF and transfer into a 15 ml falcon. To each well of a 24-well plate, thrombin (12.5 µL, 50 U/ml, Sigma-Aldrich) was added, followed by 500 µL of the cell suspension directly onto the thrombin, mixing well to give a homogenous clot which was incubated with 1 ml of EGM-2 media per well for 72 hours. Individual beads were imaged using 10X magnification with an Olympus Fluorescence Microscope equipped with a camera (25 beads per conditions) and ImageJ used to measure mean sprout length and average sprout number per bead-10 beads per treatment condition, across 4/5 individual sample wells.

### Click-iT Plus EdU Cell Proliferation Assay

Quantification of cell proliferation was performed using the Click-iT Plus EdU cell proliferation assay, as per manufacturer’s protocol with minor modifications. HUVECs were seeded at a density of 20,000 cells per well in a 24-well plate and immersed in 1 mL ECGM-2, containing 100 μM **HI2** or vehicle control (DMSO) and incubated at 37 °C and 5% CO_2_.for 24-hours. A 1 μL aliquot of 5-ethynyl-2’-deoxyuridine (EdU-1:1000) in ECGM-2 was then added to each well and incubated for a further 2 hours. Between each addition, cells were washed twice in DPBS. Cells were fixed in 1 mL of 4% PFA for 15 minutes at room temperature. To permeabilise the cells, 1 mL of PBS containing 0.5% Triton-X100 was added, and they were further incubated for 20 minutes at room temperature. A 100 μL aliquot of the ClickIT Cocktail was added to each well and incubated for 30 minutes at room temperature, in the dark. A final application of 500 μL propidium iodide was added and incubated for 15 minutes at room temperature, in the dark. Cells were immersed in DPBS during the imaging process. Images at four distinct locations within each well were sampled using the IncuCyte® ZOOM (Sartorius) system and its proprietary ZOOM 2016A software. Images were analyzed using NIH ImageJ multi-point tool function and exported to Microsoft Excel for further data transformation.

### Hybrid Homology Model Generation

The homology model of the hybrid ectodomain was based on the IR ectodomain dimer described by Croll et al (Croll et al. 2016) which is a valid protein to model due to the high homology between IR and IGF1R (∼70% overall sequence identity). The IGF1R ‘monomer’ of hybrids was created using I-TASSER(Yang et al. 2015) and a monomer from the IR ectodomain dimer used as a template. The IGF1R monomer was merged with the IR monomer from the crystal structure and the resulting dimer prepared using the Maestro graphical user interface (Schrodinger, LLC, New York, NY, 2017) with the protein preparation wizard to minimise steric clashes between residues and create disulfide links between the two monomers.

### Virtual-High Throughput Screening

A diverse set of 100 000 molecules was constructed from the ZINC diversity library of drug-like molecules using Canvas (Schrodinger, LLC, New York, NY, 2017) on the ARC3 high-performance computing cluster. This library was docked to the hybrid homology model using Glide Standard precision mode on the ARC3 high-performance computing cluster (Halgren et al. 2004). The small molecules were then ranked according to predicted binding affinity (Glide SP score). The top 5000 scoring molecules were re-docked using an alternative docking software, AutoDock 4.2(Trott et al. 2010). The 5000 molecules were clustered by energy (predicted binding affinity) and an arbitrary cut-off of -7.5 was chosen for visually inspection. Compounds that satisfied these criteria were cross-referenced with the top ranking molecules from Glide docking to identify those which were predicted to bind favourably in both docking softwares. We envisioned that a consensus docking approach (and orthogonal scoring functions) would increase our hit rate. Additional *in silico* analysis was performed using Maestro 10.2 (Maestro, Schrodinger, LLC, New York, NY, 2017) and the in house de novo design software SPROUT(Law et al. 2003) to identify compounds predicted to make hydrogen bonding and ver der waals interactions with the insulin receptor and steric clashes with the IGF1 receptor.

### Ligand-Based Virtual Screening

A shape similarity screen was performed using OpenEye’s ROCS software (OpenEye Scientific Software, Santa Fe, NM. http://www.eyesopen.com)(Hawkins et al. 2007, Hawkins et al. 2010). ROCS allows rapid identification of active compounds by shape comparison, through aligning and scoring a small molecule database against a query structure. EON compares electrostatic potential maps of pre-aligned molecules to a query structure, producing a Tanimoto similarity score for comparison. Therefore, EON can be used to analyse output structures prealigned using ROCS analysis, identifying molecules with similar similarity and electrostatics to an input structure. The **HI1** query file was the docked pose from the initial virtual screening run and the library used was the University of Leeds’ MCCB library of low energy conformers generated by OMEGA. The library contains around 27,000 molecules. The actual compounds are stored at 14 °C in 100% DMSO stock solutions, within 0% humidity cabinets. Standard parameters were used as well as an EON input command. The top 1000 ranked molecules were taken into EON (electrostatic matching) and the top 100 molecules visually inspected using VIDA. Molecules were selected for biological evaluation based on favourable shape and electrostatic matching to **HI1**.

### Statistics

Results are expressed as mean±SEM. Comparisons within groups were made using paired Student t tests and between groups using unpaired Student t tests, as appropriate. P<0.05 was considered statistically significant.

## Data Availability

This study includes no data deposited in external repositories.

## Acknowledgements

The authors would like to thank Dr Tarik Issad (INSERM Institute, Paris) for the gift of IR-Rluc and IGF1R-YPET plasmids. The authors acknowledge Dr Christopher M. Pask for small molecule X ray crystallography experiments. This work was supported by the British Heart Foundation-grant number RG/15/7/31521 and CH/13/1/30086.

## Author Contributions

MK and KJS designed and directed the study. KJS carried out the molecular modelling and compound selection. DJB and CWGF contributed to the study design. CM, HV, NH and KB carried out and analysed western blotting data. KS, ST, LL and MM synthesised HI analogues. PJM and EC carried out and analysed bead sprouting assays. All authors analysed the data and discussed the results. MK, KJS and CW prepared the manuscript.

## The Paper Explained

### Problem

A single receptor coupling nutrition to cellular growth and metabolism remains important in less complex animals. During evolution this single receptor developed into a complex system of cell surface receptors specialized to regulate metabolism (the IR) and growth (the IGF1R). A third receptor IR/IGF1R hybrid is expressed as abundantly as IR and IGF1R but unlike IR and IGF1R the role of hybrids has remained a mystery.

### Results

We describe here the discovery, synthesis and application of a small molecule allowing the first demonstration of the effect of selective reduction of hybrids in human cells. We show that reducing hybrid formation in endothelial cells impacts on insulin signalling leading to changes in EC function distinct from published reports of the effect of reducing either IR or IGF1R expression.

### Impact

Our small molecule may be used in future studies as a tool compound to investigate the complex biology of the insulin/IGF1 signalling system at a tissue, organ, and whole organism level.

